# Rituximab treatment for IgA vasculitis: A systematic review

**DOI:** 10.1101/676908

**Authors:** Cristina Carbonell, Jose-A Mirón-Canelo, Sandra Diez-Ruiz, Miguel Marcos M, Antonio-J Chamorro

## Abstract

**Objetives:** Immunoglobulin A vasculitis (IgAV) is an inflammatory disease with a controversial treatment based in corticosteroids as first line. To review cases of patients with IgA vasculitis (IgAV) treated with rituximab (RTX) and assess disease characteristics, treatment efficacy and safety.

**Methods:** We conducted a systematic literature review according to PRISMA guidelines. We searched Pubmed, Web of Science, Embase and Scopus, selecting articles with information on IgAV and RTX treatment up to February 2019, with no language limitations. We extracted data on patient characteristics, disease evolution, treatment safety and efficacy. We created a database and analyzed it using statistical software package SPSS v 22.0.

**Results:** We extracted clinical data for 43 IgAV patients treated with RTX. Distribution by sex was similar, and the median age at diagnosis was 16 (range 2 months to 70 years). The majority of patients were diagnosed at a pediatric age (24 patients under 18, 55.81%). The time of disease evolution until RTX administration greatly varied. An important number of patients suffered renal complications (86%) before RTX treatment. The frequency of adverse effects from RTX was low (7%, hypersensitivity with no treatment interruption). In terms of disease evolution, 41 patients (95.3%) presented clinical improvement, 15 patients had recurrence after initial remission (34.9%) and 34 patients (79.1%) achieved full remission after completing treatment. None of the patients treated with RTX and included in our systematic review died.

**Conclusions:** RTX is efficacious in patients with IgAV. A high percentage of patients achieved complete remission with a favorable safety profile.

## Introduction

Immunoglobulin A vasculitis (IgAV), previously called Henoch-Schönlein purpura (HSP), is an inflammatory disease. It is a small-vessel leukocytoclastic vasculitis [1] and is the most frequent vasculitis occurring in infancy [2]. The ethiopathology of this disease is still not completely understood [2, 3]. In 30-50% of cases it is possible to identify a trigger factor, primarily ear, nose, throat or respiratory infections, consumption of certain medications or toxins, or the presence of tumors [4]. From a clinical perspective IgAV is a systemic disease that may affect multiple organs [4]. IgAV diagnosis is based primarily on clinical criteria supported by histopathologic findings. The current classification criteria are those proposed by the European League Against Rheumatism/Paediatric Rheumatology International Trials Organisation/Paediatric Rheumatology European Society (EULAR/PRINTO/PRES) [5], due to their high sensitivity and specificity.

Treatment is controversial. Good quality evidence is lacking to support treatment strategies in the majority of cases, particularly refractory cases, where treatment has consisted of intravenous immunoglobulin administration, plasmapheresis and rituximab (RTX). [2–4, 6, 7] RTX, an anti-CD20 chimeric monoclonal antibody, has been widely used for hematological and rheumatologic diseases, and has been successfully used in vasculitis cases involving antibody formation and immune complex deposition. RTX reduces B cell levels and therefore IgA production, helping to control the disease [2]. The most common adverse effects are hypersensitivity and immunosuppression-related infections.

Given the lack of scientific evidence to assess the response and repercussions of using RTX to treat IgAV in either pediatric or adult populations, the objective of our study was to conduct a systematic literature review of efficacy and safety of RTX treatment for IgAV.

## Materials and methods

We conducted a systematic review according to PRISMA guidelines. We searched Pubmed, Web of Science, Embase and Scopus, selecting articles with information on IgA vasculitis and rituximab treatment up to February 2019. We used the following search terms: “Henoch Schönlein”, “Schönlein Henoch” “IgA vasculitis”, “Immunoglobulin A vasculitis”, “Rituximab”. Also, by reviewing the references in the most relevant articles we identified additional articles of interest. The search was conducted by two investigators (CC and SDR) with advice and review by two supervisors (AJC and MM). No articles were discarded based on language used.

We included in our review all cases with available data for a set of epidemiological, clinical, diagnostic and therapeutic variables (see S1 file: supplementary material).

### Statistical Analysis

We generated a database of the extracted data and analyzed it using SPSS v 22.0 statistical package. We described the distribution of continuous variables as the number of observations, either average and standard deviation or median and range. We presented the qualitative variables as the number and percentage of patients in the most relevant variables. We used McNemar’s test to compare the trends between groups of qualitative variables, with a significance level of alpha = 0.05.

## Results

Our search up to February 2019 yielded 161 articles based on our criteria, of which 139 were discarded for reasons described in Fig 1. Of the 22 articles selected (see S1 file: supplementary material) clinical data was extracted for 43 patients treated with rituximab that met the 1990 American College of Rheumatology (ACR) 1990 IgAV classification criteria [8] or the 2010 EULAR/PRINTO/PRES classification criteria [5].

**Fig 1:**
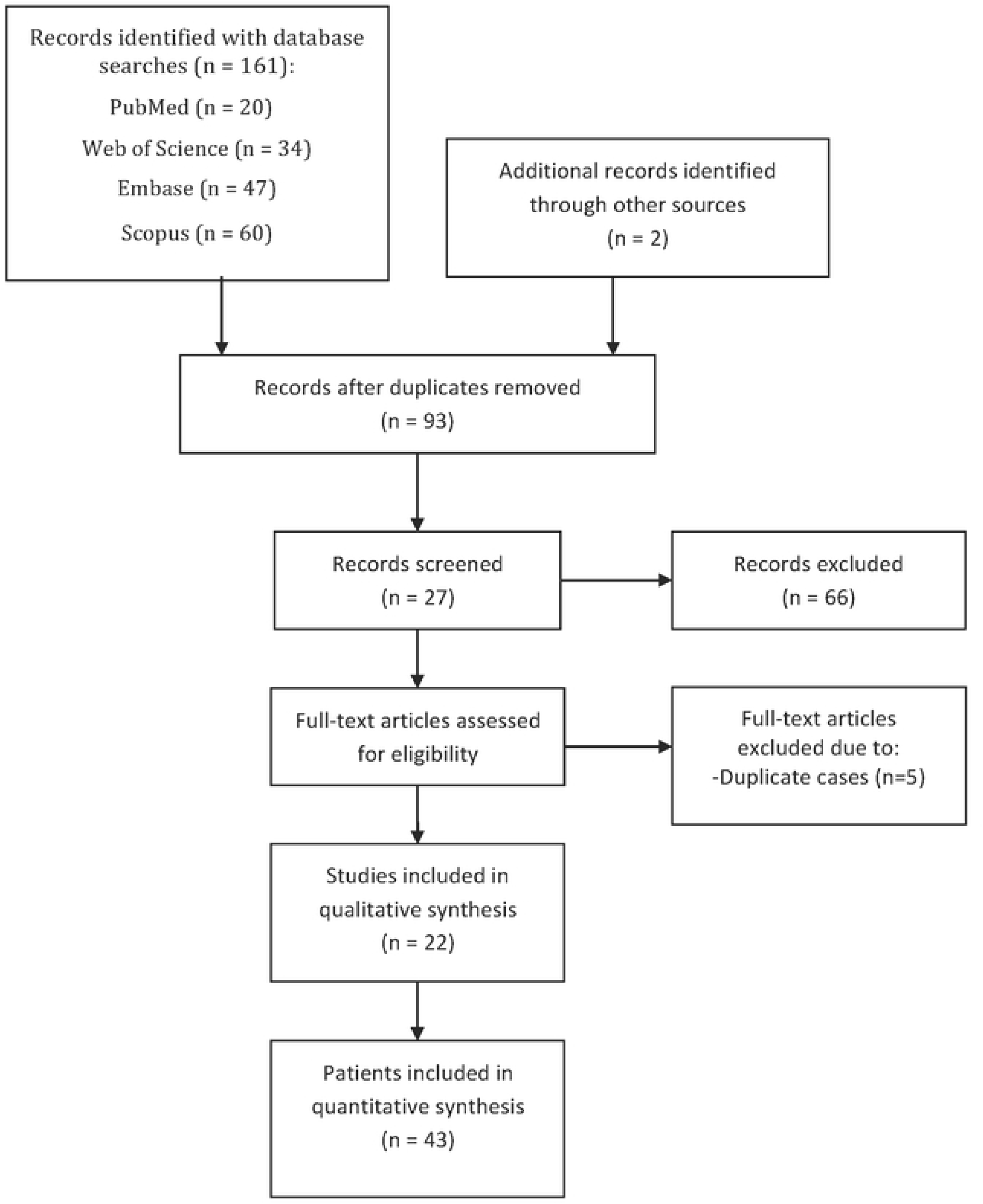
Flowchart of the selection of studies for inclusion in our systematic review.

We observed a sex distribution of 21 females (48.8%) and 22 males (51.2%), an average patient age of 22.93 years and a median age of 16 years (range: 2 months to 70 years). The majority of patients were diagnosed at pediatric age (24 patients younger than 18, 55.81%). As for ethnicity, 25 were Caucasian (58.1%), 2 were Asian (4.7%) and 2 were Hispanic. For 14 patients there was no explicit ethnicity information.

Time of disease evolution until administration of RTX varied from less than a year in 34 patients (79.06%) to 1-10 years in 6 patients (13.95%) and greater than 10 years in 2 patients (4.65%). In one patient time of disease evolution was not specified.

The principal clinical, diagnostic, treatment and evolution descriptive factors are shown in Table 1. A significant number of patients suffered renal complications N (86%) while the frequency of adverse effects with RTX was quite low (7%, hypersensitivity without treatment interruption). In terms of disease evolution, 41 patients (95.3%) presented clinical improvement, 15 patients (34.9%) suffered recurrence after initial remission and 34 patients (79.1%) achieved complete remission after treatment follow-up.

**Table 1:**
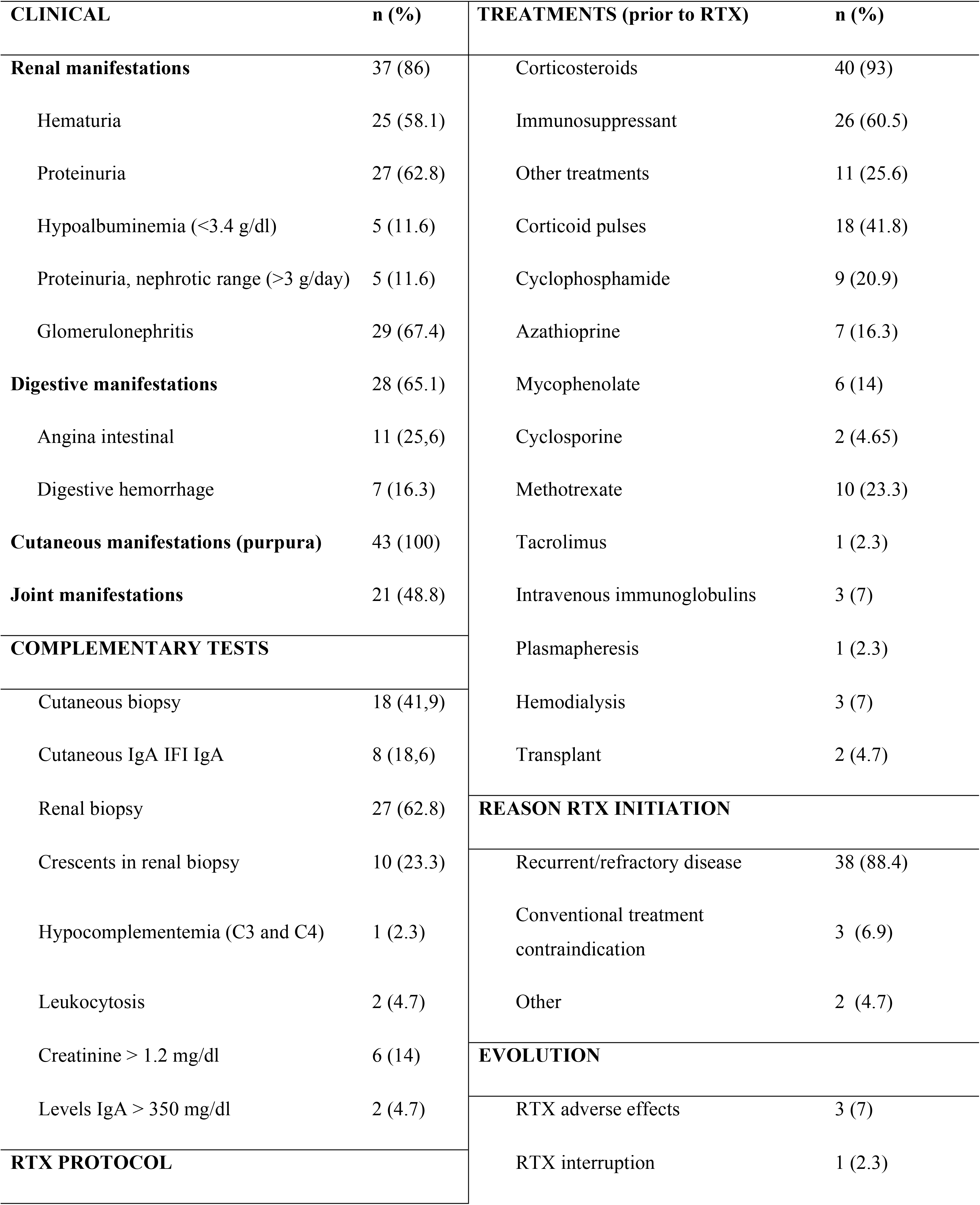
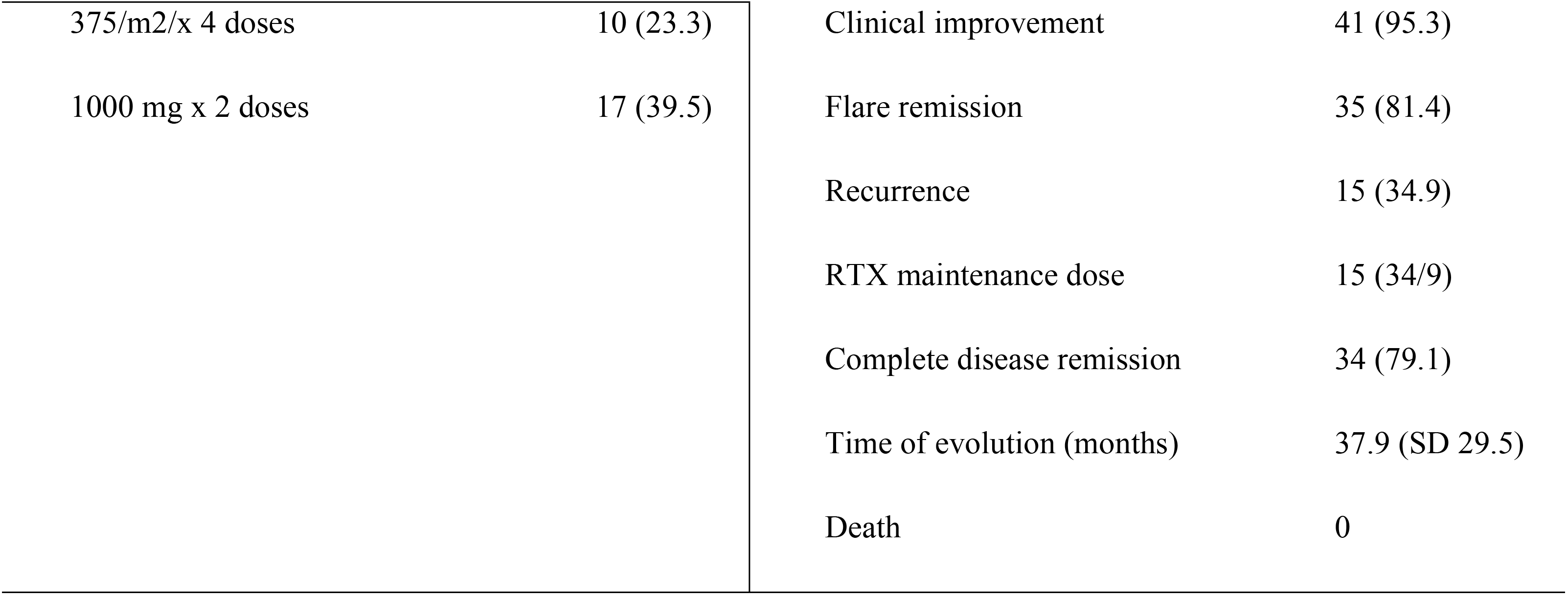
Data from patients with IgAV (n=43)

| **CLINICAL** | **n (%)** | **TREATMENTS (prior to RTX)** | **n (%)** |
| --- | --- | --- | --- |
| **Renal manifestations** | 37 (86) | Corticosteroids | 40 (93) |
| Hematuria | 25 (58.1) | Immunosuppressant | 26 (60.5) |
| Proteinuria | 27 (62.8) | Other treatments | 11 (25.6) |
| Hypoalbuminemia (<3.4 g/dl) | 5 (11.6) | Corticoid pulses | 18 (41.8) |
| Proteinuria, nephrotic range (>3 g/day) | 5 (11.6) | Cyclophosphamide | 9 (20.9) |
| Glomerulonephritis | 29 (67.4) | Azathioprine | 7 (16.3) |
| **Digestive manifestations** | 28 (65.1) | Mycophenolate | 6 (14) |
| Angina intestinal | 11 (25,6) | Cyclosporine | 2 (4.65) |
| Digestive hemorrhage | 7 (16.3) | Methotrexate | 10 (23.3) |
| **Cutaneous manifestations (purpura)** | 43 (100) | Tacrolimus | 1 (2.3) |
| **Joint manifestations** | 21 (48.8) | Intravenous immunoglobulins | 3 (7) |
| **COMPLEMENTARY TESTS** |  | Plasmapheresis | 1 (2.3) |
| Cutaneous biopsy | 18 (41,9) | Hemodialysis | 3 (7) |
| Cutaneous IgA IFI IgA | 8 (18,6) | Transplant | 2 (4.7) |
| Renal biopsy | 27 (62.8) | **REASON RTX INITIATION** |  |
| Crescents in renal biopsy | 10 (23.3) | Recurrent/refractory disease | 38 (88.4) |
| Hypocomplementemia (C3 and C4) | 1 (2.3) | Conventional treatment contraindication | 3 (6.9) |
| Leukocytosis | 2 (4.7) | Other | 2 (4.7) |
| Creatinine > 1.2 mg/dl | 6 (14) | **EVOLUTION** |  |
| Levels IgA > 350 mg/dl | 2 (4.7) | RTX adverse effects | 3 (7) |
| **RTX PROTOCOL** |  | RTX interruption | 1 (2.3) |
| 375/m2/x 4 doses | 10 (23.3) | Clinical improvement | 41 (95.3) |
| 1000 mg x 2 doses | 17 (39.5) | Flare remission | 35 (81.4) |
|  |  | Recurrence | 15 (34.9) |
|  |  | RTX maintenance dose | 15 (34/9) |
|  |  | Complete disease remission | 34 (79.1) |
|  |  | Time of evolution (months) | 37.9 (SD 29.5) |
|  |  | Death | 0 |

We evaluated the effect of RTX by analyzing CD19+ lymphocyte values before and after RTX infusion. Values before infusion were recorded in 6 of the 43 patients (13.95%) with an average of 515/mm^3^ (range: 111-1580, median: 320). Values after infusion were recorded in 14 patients (32.55%), with 11 patients showing < 1/mm^3^ and 3 patients showing > 6/mm^3^, which is considered the depletion value (average: 3.5, median: 0, range: 0-29).

We evaluated therapeutic response to RTX (before and after) from a clinical and therapeutic perspective. We saw statistically significant improvement (*p* < 0.05) in all the comparisons made (see Table 2) except for creatinine levels which did not show improvement.

**Table 2:** Evolution of clinical and therapeutic response to rituximab (before-after)

|  | Before<sup>1</sup>: n (%) | After: n (%) | P-value |
| --- | --- | --- | --- |
| **Clinical** |  |  |  |
| Cutaneous manifestations | 42 (100) | 2 (4.8) |  |
| Joint manifestations | 21 (50) | 1 (2.4) | $p < 0.001$ |
| Digestive manifestations | 34 (81) | 1 (2.4) | $p < 0.001$ |
| Renal manifestations | 36 (85.7) | 6 (14.3) | $p < 0.001$ |
| Creatinine > 1.2 mg/dl | 5 (50) | 2 (20) | NS |
| **Use of medication** |  |  |  |
| Corticosteroids | 40 (93) | 15 (34.5) | $p < 0.001$ |
| Immunosuppressants | 26 (60.5) | 10 (23.3) | $p < 0.001$ |

<sup>1</sup>Information was not available for every aspect for all patients

Results for times of remission were mixed, with an average time of 37.9 months and a range between 2 ½ months and 8 years. For 14 of the patients there was no remission information. Finally, in terms of mortality, none of the patients treated with rituximab in our systematic review died.

## Discussion

To our knowledge, ours is the first systematic literature review of IgAV vasculitis treatment with RTX in pediatric and adult populations, although we did find a narrative literature review [9] and a retrospective study of IgAV patients treated with rituximab [10].

Demographically for the 43 patients in our study, there was a slight predominance of males, which may correspond to the overall distribution of males in the adult population, and likewise a predominance of pediatric cases, where there was an even sex distribution [4]. The average age at time of diagnosis in our review was 22.93 years with a median of 16 years, which is greater than the average age of 6 years described in the bibliography for general patients with IgAV [3]. Most patients treated with RTX are primarily adult [9, 10], which may indicate that the population requiring RTX treatment showed a later onset of disease (e.g. greater frequency of renal manifestations in adult patients with IgAV vs. pediatric patients).

In our study 88.4% of the patients had experienced at least one flare prior to the flare treated with rituximab, which shows the presence of disease that was either recurrent or refractory to conventional treatment. We analyzed the presence of manifestations before and after RTX treatment. Overall, the patients analyzed presented primarily renal manifestations followed by digestive manifestations. These results are similar to other studies [10]. All manifestations apart from creatinine levels significantly improved after treatment.

In terms of diagnosis, only 18.6% of patients had histologically confirmed presence of cutaneous IgA immune complex via immunofluorescence. In the remaining patients, the presence of leukocytoclastic vasculitis after skin biopsy (41.9%) or the presence of purpuric lesions together with other symptoms were sufficient for diagnosis, based on the abovementioned classification criteria [5, 8]. The most relevant parameters were serum IgA levels, complement C3 and C4 levels and the presence of leukocytosis. Various authors have reported higher serum IgA levels in IgA vasculitis patients than in control patients [11]. In our review it was difficult to make a comparison with these results, since most of our patients had no immunoglobulin analytical data, and in the 6 patients with the data only 2 showed elevated levels. It has also been reported that complement activation by alternative pathways could be related to IgAV pathogenesis, which would be reflected by higher serum levels of these proteins in affected patients [12]. In our review, data concerning this point were collected for only 5 patients, and one case showed a reduction of serum complement levels.

In our review all patients except one had previously been treated with other immunosuppressants. The most frequent were cyclophosphamide, azathioprine and mycophenolate, corresponding to those reported in the literature [10]. Due to the frequent occurrence of spontaneous remission, IgAV treatment has habitually been focused on symptoms. More serious cases have been treated primarily with corticosteroids and immunosuppressants [13]. Until now, IgAV treatment with RTX has been limited to serious and refractory cases, normally with renal complications [2, 3, 7]. In our study the principal reasons for using RTX were recurrent and refractory cases. Additionally, RTX treatment was used in 3 cases where conventional treatment was contraindicated due to adverse effects, in one patient who received the R-CHOP protocol (rituximab, cyclophosphamide, doxorubicin, vincristine and prednisolone) and in another patient where RTX was the first line of treatment.

We found the administration protocols (RTX bolus weekly for 4 weeks with weight-adjusted dose of 375mg/m2/dose and 2 biweekly 1 g bolus treatments) and the low frequency of adverse effects (all of them mild) to be similar to those reported in the literature [10]. One of the most important findings of our study was that the vast majority of patients (95.3%) improved after initial treatment with RTX, although about a third of patients suffered disease recurrence, in line with other studies [10]. After a year, 79.1% of patients in our review presented complete remission of their disease, indicating a very favorable efficacy profile. The average time of remission in these patients was 37.9 months (±29,5). Our study also confirmed a statistically significant reduction of all clinical manifestations after treatment with RTX when compared with manifestations prior to treatment. Some of these manifestations have been described to be frequently refractory to conventional corticosteroid and immunosuppresor treatment [10]. In addition, after RTX many patients could discontinue prolonged corticosteroid and immunosuppressant treatment. Among patients in whom the disease was still active, the most frequent clinical manifestation was renal disease, in agreement with the literature, probably reflecting chronic and irreparable damage after disease flares. None of the patients in our review had died, in agreement with other studies showing minimal mortality rates [10].

The principal limitations of our study were: a) the small sample size, although we expanded our systematic search to many databases without language limitations, and introduced all possible quantifiable variables; b) the lack of cutaneous or renal biopsies with IFI for IgA for some patients, although all patients met the standard classification criteria and had compatible cutaneous manifestations; and c) the highly heterogenous information available from each of the articles in our review.

In summary we can conclude that RTX is an efficacious and safe treatment for IgAV with few side effects, and it could be a good therapeutic alternative primarily for those patients where conventional immunosuppressant treatment is contraindicated, or patients with recurrent, refractory or severe clinical course. These conclusions, and the effect of RTX on specific manifestations should be corroborated with clinical trials in adult and pediatric populations.

## PRISMA 2009 Checklist statement

The authors have read the PRISMA 2009 Checklist, and the manuscript was prepared and revised according to the PRISMA 2009 Checklist.

## Supporting Information

S1. File. 1. Supplementary material: List of variables included and list of articles selected for the systematic review.

S2. File. 2. PRISMA checklist.

